# Optimizing magnetometers arrays and analysis pipelines for multivariate pattern analysis

**DOI:** 10.1101/2023.09.21.558786

**Authors:** Yulia Bezsudnova, Andrew J. Quinn, Ole Jensen

## Abstract

**Background:** Multivariate pattern analysis (MVPA) has proven an excellent tool in cognitive neuroscience used M/EEG, and MRI. It also holds a strong promise when applied to optically-pumped magnetometer-based magnetoencephalography.

**New method:** To optimize OPM-MEG systems for MVPA experiments this study examines data from a conventional MEG magnetometer array, focusing on appropriate noise reduction techniques for magnetometers. We also determined the least required number of sensors needed for robust MVPA for image categorization experiments.

**Results:** We found that the use of signal space separation (SSS) significantly lowered the classification accuracy considering a sub-array of 102 magnetometers or a sub-array of 204 gradiometers. We also found that classification accuracy did not improve when going beyond 30 sensors irrespective of whether SSS has been applied.

**Comparison with existing methods:** The power spectra of data filtered with SSS has a substantially higher noise floor that data cleaned with SSP or HFC. Consequently, the MVPA decoding results obtained from the SSS-filtered data are significantly lower compared to all other methods employed.

**Conclusions:** When designing an MEG system based on SQUID magnetometers optimized for multivariate analysis for image categorization experiments, about 30 magnetometers are sufficient. We advise against applying SSS filters to data from MEG and OPM systems prior to performing MVPA as this method, albeit reducing low-frequency external noise contributions, also introduces an increase in broadband noise. We recommend employing noise reduction techniques that either decrease or maintain the noise floor of the data like signal-space projection, homogeneous field correction and gradient noise reduction.

**Highlights:**

- A sensor array of about 30 sensors is sufficient for multivariate pattern analysis using conventional MEG magnetometers for image classification.
- Using signal space separation filter on magnetometer data prior to multivariate pattern analysis might reduce classification accuracy due to an increase in white noise in the data contributed by the algorithm.
- When performing multivariate data analysis, other noise reduction approaches that diminish the contribution of external noise sources and reduce the variance of the data are advisable such as synthetic gradiometers, signal space projection or homogeneous field correction.

## 1. Introduction

Multivariate pattern analysis (MVPA) is an established methodology for functional MRI, MEG, and EEG. MVPA operates by classifying the spatial distribution of brain activity associated with representing different objects’ classes or experimental conditions (Haxby et al., 2001; Haynes and Rees, 2006; Pereira et al., 2009; Grootswagers et al., 2017; Carlson et al., 2019; Stokes et al., 2015; Cichy et al., 2014). If spatial patterns of brain activation successfully predict condition or stimulus category, it implies the presence of relevant information related to the experimental manipulation within the neuroimaging data. Furthermore, applying MVPA in the context of representational similarity analysis (RSA) allows to fuse the data from BOLD measures obtained from fMRI and electrophysiological data obtained from EEG/MEG, leading to an enhanced understanding of the brain’s spatiotemporal properties in relation to neuronal responses (Kriegeskorte and Kievit, 2013; Guggenmos et al., 2018; Kriegeskorte et al., 2008; Kriegeskorte, 2009; Diedrichsen and Kriegeskorte, 2017). Overall, MVPA applied for classification approaches and RSA is a powerful tool for cognitive neuroscience widely used to study object recognition (Cichy et al., 2014; Proklova et al., 2016; Cichy et al., 2016; Kaplan et al., 2015; Dirani and Pylkkänen, 2023) or other cognitive brain functions like prediction, spatial navigation, memory and emotions (Frisby et al., 2023; Harrison and Tong, 2009; Kornysheva et al., 2019; Brown et al., 2016; Ariani et al., 2022; King and Dehaene, 2014; Wang et al., 2018).

This study lays the groundwork for exploring the potential benefits of utilizing on-scalp optically pumped magnetometer-based magnetoencephalography (OPM-MEG) in experiments designed for MVPA. There are two main reasons why MVPA stands to benefit from OPM-MEG systems. First, OPMs can be put directly on the subject head and therefore the magnitude of the MEG signal is stronger compared to the signal obtained by SQUID-MEG systems (Brookes et al., 2022; Bezsudnova et al., 2022). Second, the sensors can be arranged in an array that better captures the spatial characteristics of the brain signal (Beltrachini et al., 2021; Iivanainen et al., 2017; Brookes et al., 2021). As a result, OPM-MEG potentially has higher sensitivity and better spatial resolution than conventional MEG systems, which is beneficial for MVPA (Labyt et al., 2022; Wens, 2023). While OPM-MEG is still in its early stages of development and specific engineering challenges have to be solved, a study conducted by (Nugent et al., 2022) demonstrates the theoretical capability of a dense OPM array to effectively resolve multiple independent electrophysiological sources to a degree currently achievable only using an invasive technique.

Notably, conventional MEG studies typically involve the analysis of gradiometers due to their lower sensitivity to external noise sources (Vrba and Robinson, 2001); however, current OPM-MEG systems utilize only magnetometers (Rea et al., 2022; Alem et al., 2023; Kowalczyk et al., 2021; Gutteling et al., 2023; Hunter, 2019). Therefore, to pave the way for optimizing MVPA used with OPM-MEG, this study focuses on data obtained from a conventional MEG magnetometer array. Specifically, we explore appropriate noise reduction techniques for magnetometers as well as attempt to estimate the number of sensors required for robust MVPA.

## 2. Methods

### 2.1 Experimental paradigm

In this study, we investigated the time course of the emergence of semantic representations using picture stimuli employing MVPA applied to conventional SQUID MEG data.

Thirty-eight healthy native English adult participants took part in the study. Five participants were excluded due to excessive noise in the data, two participants were excluded due to poor performance (>15% wrong answer), therefore the final data set consists of recordings from thirty-one participants (mean age ± SD = 21.77 ± 3.31; 21 female). The study was conducted at the Centre for Human Brain Health. The University of Birmingham Ethics Committee approved the study. Participants were compensated with credits or a monetary reward.

The stimulus set was composed of 48 objects. The objects were organized as in the study (Iamshchinina et al., 2022) according to three types of categories, each divided into two groups: size (big/small), movement (still/moving), and nature (natural/man-made). Each object in the study was part of one of the sub-categories(Fig. 1 e.g. a book is small, still, and man-made). The stimulus set was balanced such that each categorical division included one-half of the stimulus set (24 objects). Hence each set of sub-categories (e.g. big, still, natural) included 6 objects. The selection of category divisions in this study was based on previous findings demonstrating reliable neural representations of semantic dimensions across categories (Huth et al., 2012; Konkle and Caramazza, 2013). For other purposes not to be analysed here, each object is presented in two modalities: written words and pictures. The size of the images was 400 × 400 pixels. Prior to the object presentation, the fixation cross was shown for 0.5 – 0.7 s. Each object was presented for 0.6 s. The experiment comprised 9 blocks, with 4 blocks where pictures were successively presented (“Picture” blocks in Fig. 1B). Only data from the picture blocks was used in this study. After each block, a participant had a break. Each stimulus was present 4 times in a block. We used a two-alternative forced choice task (question trial), a participant to identify the written word that corresponded to the immediately preceding picture stimulus during the picture blocks. The study was designed to engage participants and encourage deep processing of stimuli in a cross-modal manner. The design is illustrated in Fig. 1.

**Figure 1:**
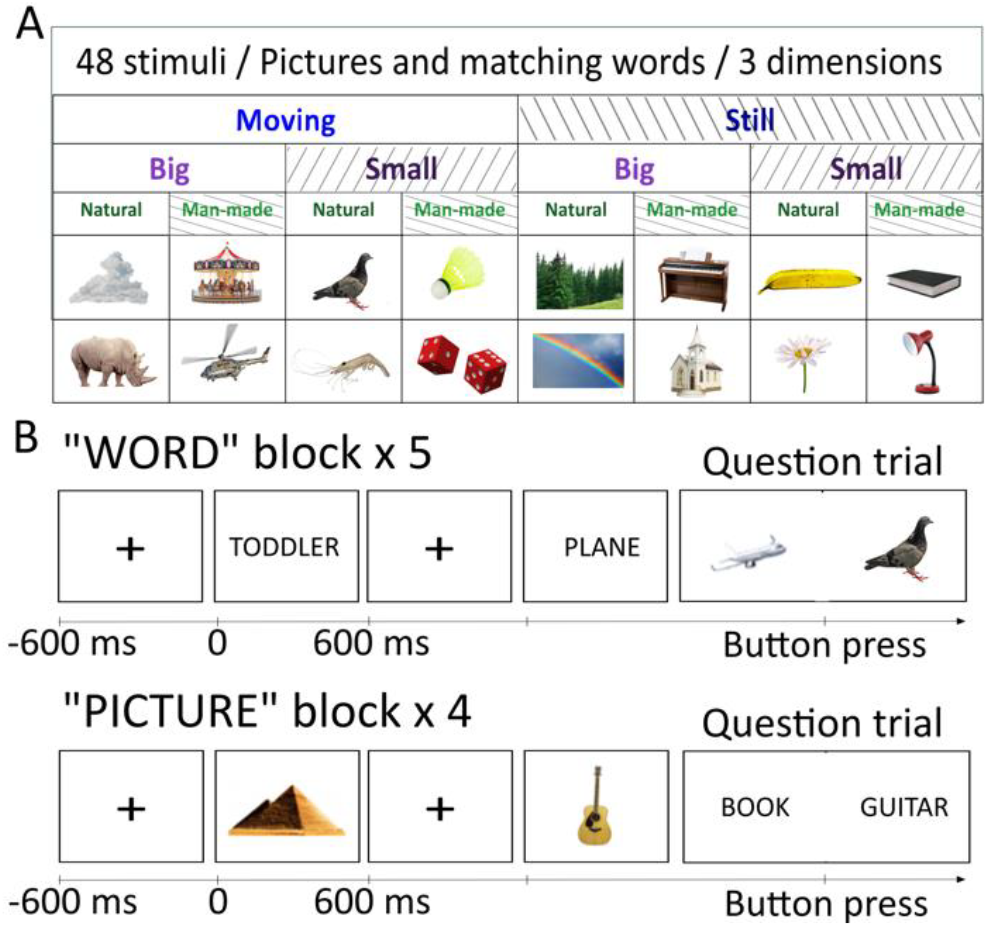
Experimental design. A) The stimulus set is comprised of 48 objects which can be divided according to 3 dimensions (“size”: big/small; “movement”: still/moving; “nature”: natural/man-made). B) In both image and word blocks, participants were presented with stimuli in random order. In the picture block, participants viewed images of the objects, while in the word block, they read the corresponding words. The task (question trial) was presented randomly every fifth trial on average. Participants were required to identify whether the picture or word corresponded to a previously seen stimulus and press the appropriate button accordingly. When pictures were presented the probe question was presented as a word and vice versa.

In each block, a randomly selected stimulus was presented on a grey screen at a visual angle of 6°. The stimuli presentation is implemented in Matlab using the Psychophysics Toolbox (Brainard and Vision, 1997). For a more detailed description of the experimental paradigm, please refer (Bezsudnova and Jensen, 2023).

### 2.2 MEG recordings

MEG data were recorded using a 306-sensor TRIUX Elekta Neuromag system, consisting of 204 orthogonal planar gradiometers and 102 magnetometers, with online band-pass filtering from 0.1 to 330 Hz and a sampling rate of 1000 Hz. Prior to data acquisition, the locations of three bony fiducial points (nasion, left and right preauricular points) were digitized using a Polhemus Fastrack system. Additionally, four head-position indicator coils (HPI coils) were digitized: two on the left and right mastoid bone and two on the forehead, spaced at least 3 cm apart. To facilitate the spatial co-registration of MEG source analysis with individual structural MRI images, a minimum of 400 extra scalp points were acquired for each participant. Following these preparations, participants were positioned upright with a 60° angle backrest under the MEG gantry. To record the horizontal EOG, a pair of electrodes was attached approximately 2.5 cm away from the outer canthus of each eye. To record the vertical EOG, a pair of electrodes was placed above and below the right eye in line with the pupil. The ECG was recorded from a pair of electrodes placed on the left and right collarbone.

### 2.3 Standard data processing pipeline

The data was analysed using the open-source toolbox MNE Python v1.6.1 (Gramfort et al., 2013) following the standards defined in the FLUX Pipeline (Ferrante et al., 2022).

The standard MVPA pipeline for a conventional MEG consists of the following steps.

First, noisy sensors were found and were annotated. This step was implemented using “mne.preprocessing.find_bad_channels_maxwell” function. Following this, the data from the sub-array of interest, whether only magnetometers, only gradiometers, or both, are extracted.

As a next step signal-space separation (SSS) is applied to reduce magnetic artefacts from the environment.

The SSS method models a magnetic signal measured by an array of sensors using multipole expansion of the magnetic field. A signal is represented as two truncated multipole series: interior (∼1/*r*) and exterior (∼*r*), which differentiate between sources originating inside the helmet and external magnetic sources respectively. *r* is the first spherical coordinate. The SSS-filtered signal is reconstructed only from the brain-related components e.g., the truncated interior expansion, all external origin components are dropped. The truncation numbers *L*_*in*_ and *L*_*out*_ define how many terms of the interior and exterior expansion are used in the model and, therefore, define the order of spatial frequency components of the reconstructed signal (Taulu and Kajola, 2005; Taulu and Simola, 2006). This results in a substantial rank reduction of the data, typically from the number of MEG sensors to a significantly lower number (from about 306 to about 65 dimensions for a typical MEGIN system). Importantly the SSS procedure attenuates high-spatial frequency components of the signal that are not included in the truncated expansion.

It is important to note that SSS is implemented as an oblique projection: the basis of the interior and exterior expansions are uncorrelated but not orthogonal. Therefore, when reconstructing the signal using the interior basis, this may result in white noise being amplified (Taulu and Kajola, 2005; Tierney et al., 2023). However, for the MEGIN system with 306 sensors, typically the reduction of external interference outweighs the increase in white noise, resulting in an overall lower noise level after applying the SSS filtering. This is particularly the case for lower frequencies.

As mentioned in the introduction, this study is focused on finding optimal noise-reduction techniques for magnetometers. The alternative method to SSS filtering developed for OPM data is homogeneous field correction (HFC) (Tierney et al., 2021, 2022). HFC uses the same multipole expansion of magnetic field as the SSS method to model a measured signal. However, HFC projects out the exterior part of the expansion as opposed to reconstructing a signal from the interior part as done by SSS. Orthogonalizing the data with respect to the modelled interference (exterior expansion) ensures that the variance of the filtered data will be lower than that of the original data. However, this approach may suppress some brain-originated signal that happens to be correlated with the modelled noise (Tierney et al., 2023). HFC only reduces the rank of the data by about 2-3 components.

Another common method to reduce environmental noise is the signal-space projection (SSP) technique (Uusitalo and Ilmoniemi, 1997). SSP on the assumption that the magnetic field distributions generated by the sources in the brain have spatial distributions sufficiently different from those generated by external noise sources. SSP achieves noise reduction by projecting the data onto a subspace that is orthogonal to the spatial patterns associated with noise, known as projectors. These spatial projectors are created from the vectors that remove the most prominent signals from the noisy empty room recordings using principal component analysis. Note that SSP may also suppress some brain-originated signals that spatially correlated with the projectors. This procedure typically reduces the rank of the data by about 8 components.

In this work, we omitted the step of employing independent component analysis typically used to remove components associated with cardiac artefacts and eyeblinks (Hyvärinen and Oja, 2000)), in order to assess the impact of different filtering techniques on classification accuracy. As a next step, the data was low-pass filtered at 100 Hz to reduce HPI coils artefacts. Following this, the sensors flagged as “bad” were excluded from further processing, and remaining data was segmented into trials. Trials corresponding to the same stimulus were averaged together to construct “supertrials”. This step increases the signal-to-noise of the data (Ashton et al., 2022; Guggenmos et al., 2018).

Prior to MVPA, we downsampled the data to 500 Hz. Epochs are cropped to 0.1s before stimuli onset and 0.7s after. Each time point was represented as a 306-dimensional vector (sensor data). We expanded this feature by including past 25 ms (12 time points) and future 25 ms (12 time points) of data, resulting in 306×25 dimensional feature vector representing each time point. This procedure (termed “delayed embedding”) resulted in a more information-rich representation of the neural activity associated with a given stimulus by incorporating more time points (Chan et al., 2011; Tyler et al., 2013; Cheng, 2021). This feature vector was then used in the classifier.

MVPA was applied to the pre-processed MEG data to classify the categories of the presented objects. Prior to classification, the data was standardized by removing the mean and scaling to unit variance per sensor and detrended. Subsequently, we employed a support vector machine (SVM) (Cortes and Vapnik, 1995) from the Python module Scikit-learn (Pedregosa et al., 2011) to classify the data over time. The classification procedure relied on a 5-fold cross-validation approach, and the performance was quantified using the area under the curve (AUC) metric, which measures the classifier discriminative ability. We computed the decoding curve for each category dimension separately and subsequently averaged them.

Throughout this study, we used the following annotations to present the results. The pipeline incorporating the SSS filter is denoted as “SSS”. Unless otherwise specified, we employed standard truncation values of *L*_*in*_ = 8 and *L*_*out*_ = 3. The “HFC” pipeline denotes the noise attenuation using the HFC filter. In the case of the HFC filter, we utilized second-order correction, which accounts for both homogeneous field components and the first-order gradients. The pipeline labelled “SSP” employed the SSP algorithm for noise reduction. We used system-provided projectors, e.g. projectors calculated from the empty-room recording prior to the experiment (not on the same day). The pipeline labelled as “raw” did not involve any noise reduction algorithms aside bad sensors rejection which was also performed for every other pipeline.

### 2.4 Number of channels

We asked how many magnetometers are needed to get good classification accuracy results. Analysing how classification accuracy scales with the number of sensors in the array would provide insight into the spatial frequency of the task-relevant brain signal. We randomly selected *N* magnetometers, spaced equidistantly, from the existing pool of 102 sensors. *N* was varied from 7 to 102 with step 5. The data from the chosen set of sensors were used in a feature vector, which included 25 ms of past and future time points. This feature vector was subsequently applied to train and test the classifier. Note that this analysis was done only for the “SSP” and “HFC” pipelines.

### 2.5 Spatial distribution of the task-relevant brain signal

To further examine the spatial distribution of the brain signal underlining the classification of categories we computed the topographical activation patterns back-projected from the classifier weights using the method described in Haufe et al. (2014). We used SSP-filtered data.

### 3.6. Statistical analysis

To identify the significant time points for the classification, we used a non-parametric one-sampled permutation t-test (one-tailed) against 50% chance level, controlled for multiple comparisons (Sassenhagen and Draschkow, 2019; Maris and Oostenveld, 2007) implemented in the GLMTools Python package (https://pypi.org/project/glmtools/). Clusters of t-values (corresponds to an alpha threshold of *P>* 0.05) that exceeded the cluster-forming threshold (at least two time-adjacent points) were included in the permutation distribution using the sum of t-values within the cluster. The null distribution was formed by computing the largest cluster from each of the 1000 permutations, which were generated by randomly flipping the signs of the data points. A cluster was considered significant if its cluster stat (sum of t-value with cluster) was in the 95th percentile of the null distribution. This corresponds to a threshold of *P* = 0.05.

The same analysis was conducted to identify time points where two classification curves significantly differed from each other, with the difference between the two curves compared against chance level.

## 3. Results

We aimed to examine appropriate noise reduction methods for magnetometers and determine the number of sensors required to achieve robust MVPA performance. This investigation constrains design considerations for future developments utilizing MVPA applied to data from OPM-MEG systems.

The study is based on conventional SQUID-MEG data recorded when participants were looking at pictures of objects from different categories: size (big/small), movement (still/moving), nature (natural/man-made; see Fig. 1). We compared MVPA classification accuracy curves for different pre-processing pipelines utilizing SSS-, HFC-, and SSP-filtering compared to no filtering at all.

We started by examining the power spectral density of magnetometer data after each filtering method in Fig. 2. The power spectrum was calculated for the experimental data recorded while a participant was performing the task (Fig. 2A) and for empty room recordings on a different day (Fig. 2B) within a 0 - 60 Hz interval. In both datasets, there is a noticeable trend where the SSS pipeline results in a visible increase in power at frequencies above 40 Hz, while effectively reducing prominent environmental interferences like 50 Hz mains and low-frequency noise. The HFC and SSP methods result in the strongest reduction in power also at higher frequencies. However, HFC exhibits stronger attenuation of external interference, particularly noticeable in the suppression of the 50 Hz noise in Fig. 2. Additionally, HFC effectively reduces interference from the empty room recording in the 20-35 Hz range (Fig. 2B), whereas SSP fails to do so because the system-provided SSP projectors did not account for these artefacts.

**Figure 2:**
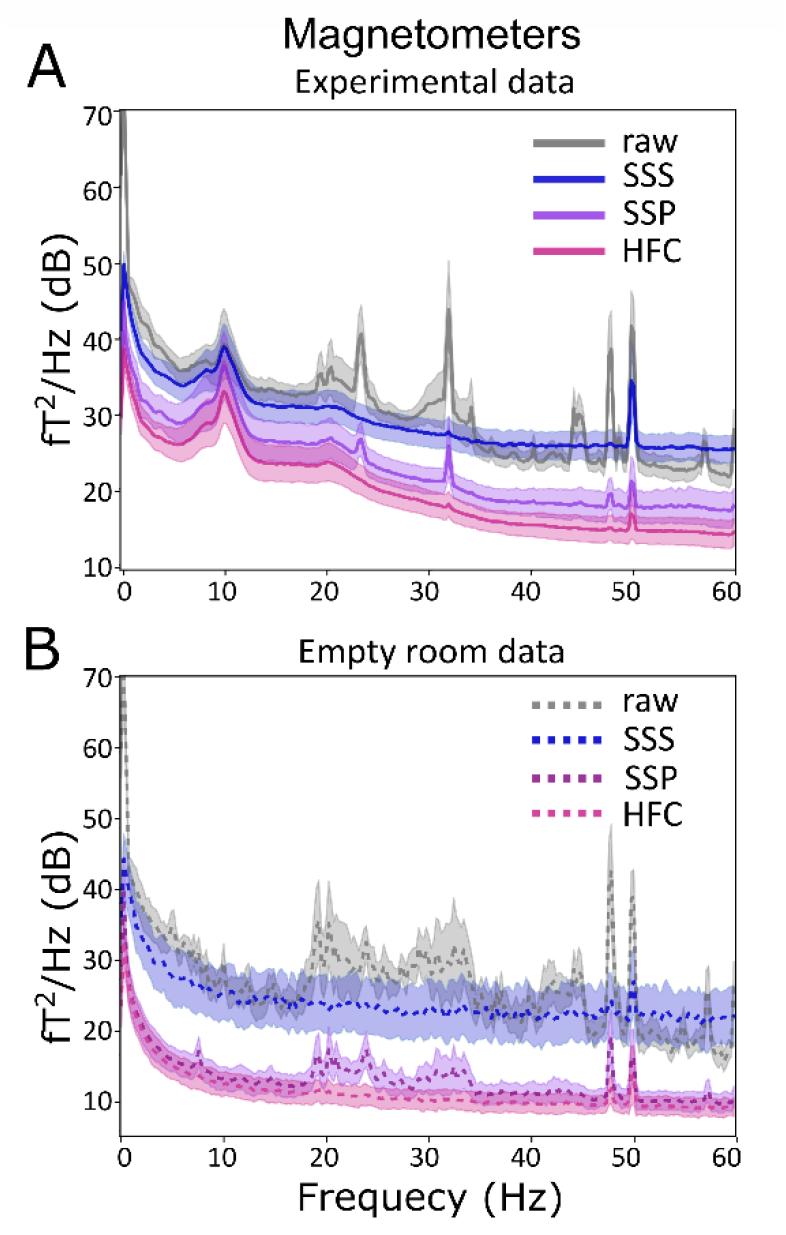
Power spectra of the magnetometer data A) of experimental data recorded while a participant was performing the task and B) of an empty room recording on a different day. The solid lines represent the averaged power spectrum, while the shaded regions denote the standard deviation interval. Grey lines: the spectra before the application of noise reduction algorithms (“raw” pipeline). Blue lines: after the application of the SSS filter (SSS pipeline). Magenta lines: after the application of HFC (HFC pipeline). Purple lines: after the application of SSP (SSP pipeline).

Overall, the data filtered with the HFC method demonstrates the lowest noise floor and the most effective attenuation of external interferences in both the experimental data and empty room recordings.

### 3.1 SSS filter does not benefit MVPA

Fig 3A shows the MVPA classification results as a function of time for the different pre-processing pipelines. We find that the use of use significantly decreases the classification accuracy considering the 102 magnetometers compared to the decoding results from the “SSP” and “HFC” pipelines. Moreover, its use does not significantly enhance the MVPA results compared to the “raw” pipeline.

**Figure 3:**
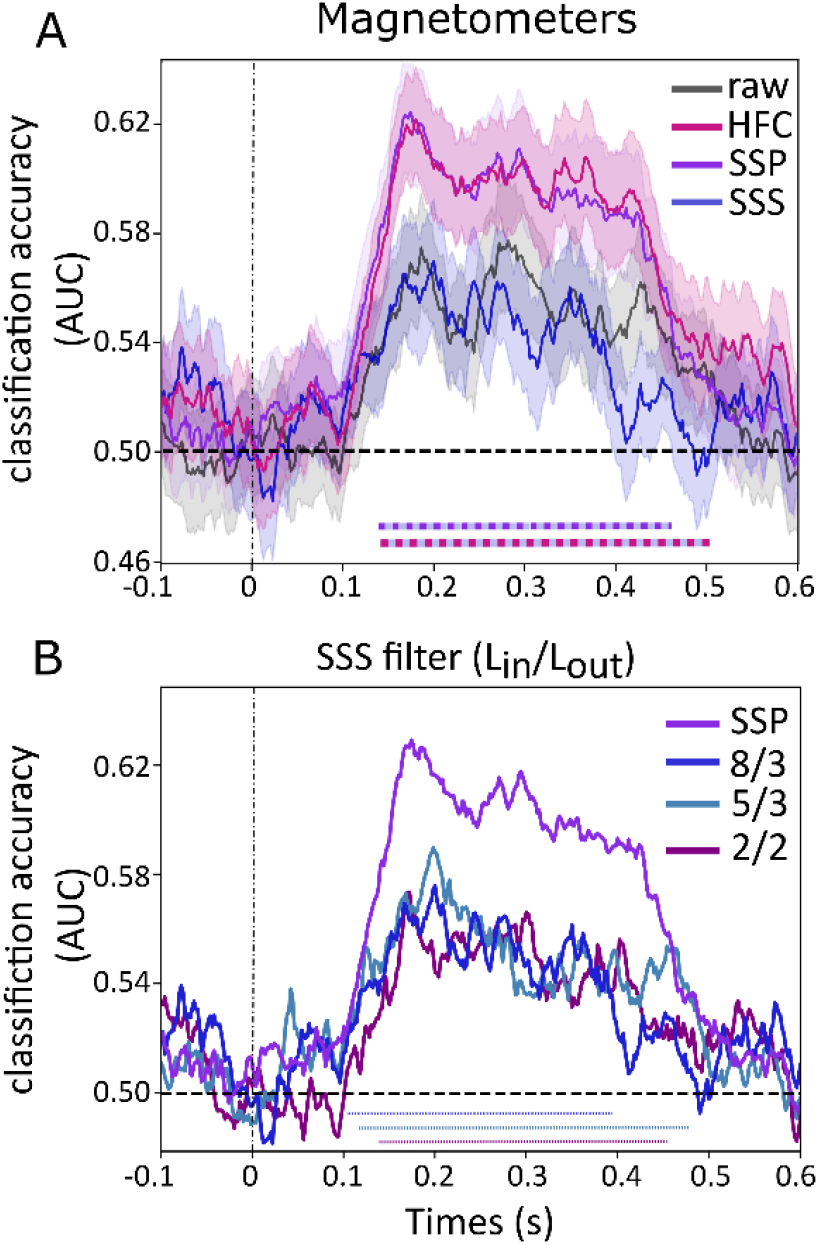
The time course of object categorization during image presentation for 102 magnetometers array. A) Comparison of the different pre-processing pipelines. Black line: “raw” pipeline. Blue line: “SSS” pipeline (*L*_*in*_ = 8, *L*_*out*_ = 3). Magenta lines: “HFC” pipeline. Purple lines: “SSP” pipeline. The shaded area reflects the standard error. Dotted line indicate time intervals where there is a significant difference (p<0.05; controlled for multiple comparisons over time) between different pipelines. Purple-Blue is between the SSS and the SSP pipelines. Magenta-Blue is between the SSS and HFC pipelines. B) Comparison of the decoding after the SSS filter with different truncation values. The coloured lines mark SSS filtered data with *L*_*in*_ = 8, *L*_*out*_ = 3; *L*_*in*_ = 5, *L*_*out*_ = 3; *L*_*in*_ = 2, *L*_*out*_ = 2. Purple line marks “SSP” pipeline. Coloured dotted lines annotate significant time of the same coloured decoding curve against the chance level. Dashed line signifies the chance level.

To examine how the SSS filter affects the MVPA results, we applied SSS filter with the different truncation values. Specifically, in Fig. 3B we show the classification curves for the SSS filter with *L*_*in*_ = 8, 5 while *L*_*out*_ = 3. We also used an extreme case of SSS filter with only the first two terms of the internal and external multipole expansions (*L*_*in*_ = 2, *L*_*out*_ = 2). Fig. 3B demonstrates that even in this extreme case the truncation value only modestly influences how the SSS filter affects the classification curve. The calculation with *L*_*in*_ = 8, 7, 5 and *L*_*out*_ = 2 produce classification curves with similar shapes, with maximum classification accuracy around 0.58, but are not shown here for clarity. *L*_*in*_ = 9 is not feasible for an array of 102 sensors especially when approximately 4 to 5 sensors are identified as noisy.

The application of the SSS filter also significantly reduces the MVPA classification accuracy when selecting a 204-gradiometer array at 0.23 – 0.41 s interval after stimulus onset (Fig. 4A). On the contrary, the application of the SSS filter does not lead to a significant decrease in MVPA classification when all 306 sensors are used (Fig. 4B).

**Figure 4:**
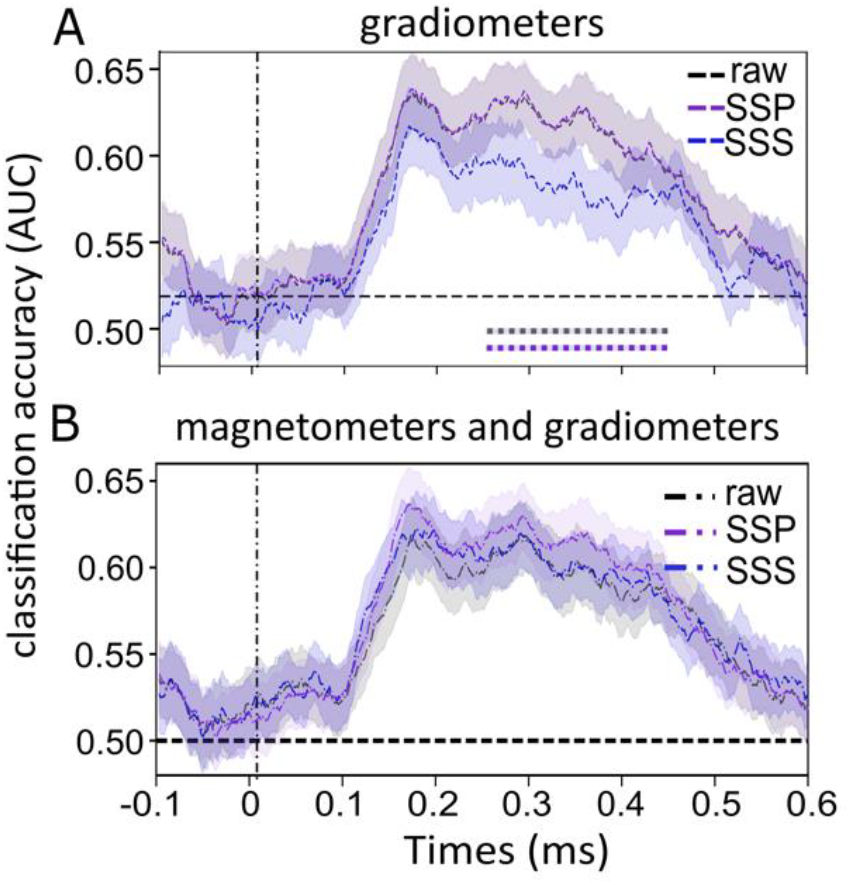
The time course of object categorization during image presentation. Black line: “raw” pipeline (unfiltered data). Blue line: the “SSS” pipeline with (*L*_*in*_ = 8, *L*_*out*_ = 3). Purple line: “SSP” pipeline. A) The gradiometers data only. Dotted lines indicate time intervals where there is a significant difference (p<0.05; controlled for multiple comparisons over time) between pipelines. Purple and blue is between “SSP” and “SSS”. Purple and blue is between “raw” and “SSS”. Note that “raw” and “SSP” lines overlap B) The magnetometer and gradiometer data combined. The shaded area reflects the standard error, while a dashed line signifies the chance level.

### 3.2 Magnetometers and Gradiometers

After optimizing the pre-processing pipeline, we compared the performance of 102 magnetometers and a subset of 102 gradiometers using only “raw” and “SSP” pipelines, as the HFC filters were developed specifically for magnetometers. On each magnetometer location, one of two planar gradiometers was selected randomly. The results shown in Fig. 5A demonstrate that the gradiometer array performs significantly better than the magnetometer array during the 0.13 – 0.27 s and 0.33 – 0.41 s interval when using the “raw” pipeline. The use of the gradiometer array yields higher classification accuracy due to the sensors’ intrinsic environmental noise suppression. However, the SSP-filtered data from both magnetometer and gradiometer arrays do not exhibit significant differences (Fig. 5B). Therefore, for robust MVPA performance, utilizing gradiometers (whether intrinsic (Cook et al., 2024) or synthetic (Vrba and Robinson, 2001) is as effective as denoising magnetometer data with SSP or HFC filters.

**Figure 5:**
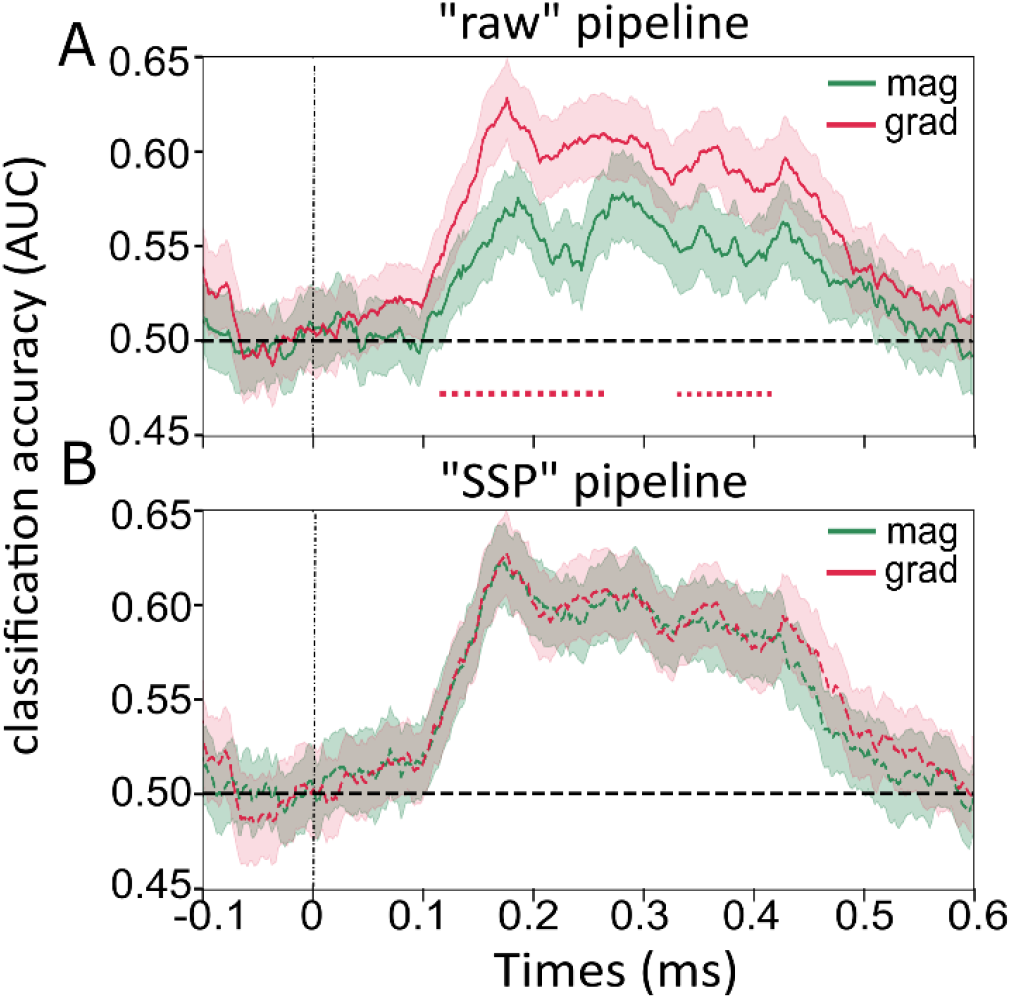
The time course of object categorization during image presentation for A) “raw” pipeline and B) “SSP” pipeline. Green line is for 102 magnetometers array. Red line is 102 gradiometers array. The shaded area reflects the standard error, while a dashed line signifies the chance level.

### 3.3 Number of sensors

Finally, we investigated the impact of the number of sensors on MVPA classification accuracy. To characterize the decoding curve for each subset of sensors we calculated the area under that decoding curve (0.15 - 0.45 s) averaged across participants. This time was chosen to capture when the brain processes and encodes relevant information related to the object category. Fig. 6A illustrates the relationship between AUC of classification as a function of a number of magnetometers. We observe that classification saturates at about 30 sensors for both “SSP” and “HFC” pipelines. The SSS or “raw” pipelines applied on 102 sensors array do not reach the AUC level, at which the other two pipelines saturate.

**Figure 6:**
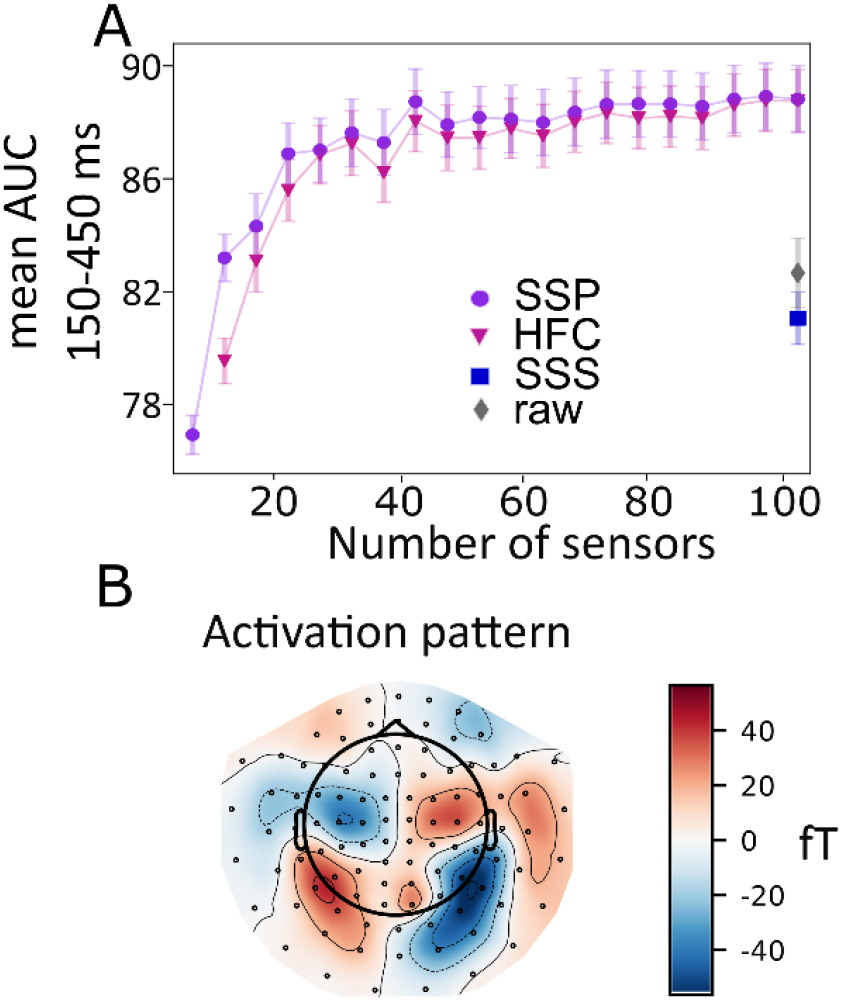
A) AUC of the decoding curve over a time interval 150-450 ms after stimulus presentation is plotted as a function of the number of sensors in the A) AUC of the decoding curve over a time interval 150-450 ms after stimulus presentation is plotted as a function of the number of sensors in the magnetometer array. Purple line represents the “SSP” pipeline. Magenta line represents the “HFC” pipeline. The blue marker represents the AUC of the decoding curve for the “SSS” pipeline with 102 magnetometers, the grey marker represents the “raw” pipeline with 102 magnetometers; shaded areas and error bars reflect the standard error B) Topographical activation pattern (Haufe et al., 2014) reconstructed from the classifier weights calculated for “SSP” pipeline with 102 magnetometers averaged in the interval.

We also examined the activation patterns calculated from classifier weights using data from the “SSP” pipeline of 102 magnetometers array. Fig. 6B shows that the topographic distribution of the neurophysiological data of the task-relevant signal does not have high-frequency spatial features, instead appearing clustered together in a few distinct regions. This observation further supports that 30 equally distributed sensors are sufficient to achieve robust classification accuracy with the current experimental paradigm.

## 4 Discussion

We explored various noise reduction techniques for conventional MEG data in the context of MVPA analysis. The results demonstrate that the use of the SSP or HFC filter for denoising magnetometry data yields a significantly higher classification accuracy compared to when using the SSS filter or raw data. We also find there are no differences in classification accuracy when comparing an SSP-filtered 102 magnetometer array to an SSP-filtered 102 gradiometer array. Moreover, we show that employing only 30 magnetometers evenly distributed over the head is sufficient to achieve classification accuracy comparable to a 102 magnetometer array when using MVPA for image categorization.

Based on our results we do not recommend using the SSS filter in OPM-MEG data when conducting MVPA. The implementation of SSS filters does not significantly increase the SVM classification accuracy compared to the unfiltered data and significantly reduce decoding when compared to SSP and HFC filtered magnetometer data (Fig. 3). Moreover, SSS results in a significant reduction of the SVM classification accuracy when considering unfiltered SQUID gradiometers (Fig. 4A). There are several possible reasons why SSS significantly decreases MVPA. The first reason might be the loss of a task-relevant brain signal due to the truncation of the multipole expansions used to model the brain signal and external interference. Due to the truncation, an unmodeled brain signal with spatial frequencies higher than defined by maximum *L*_*in*_ is omitted in the filtered data. However, by comparing decoding curves with *L*_*in*_ = 2 to *L*_*in*_ = 8, we show that excluding components of brain signals with low spatial frequencies corresponding to *L*_*in*_ = [3..7] does not negatively affect MVPA results (Fig. 3B). This observation suggests that the task-relevant brain signal has a relatively low spatial frequency distribution, and the standard truncation values of SSS filtering do not lead to its loss. The low-frequency spatial distribution of the task-relevant signal is evident from the activation pattern calculated from classifier weights (Fig. 6B), which exhibit a low-frequency spatial topology. Additionally, the fact that 30 sensors perform as well as 102 sensors (Fig. 6A) further supports this observation. Hence, the truncation of the multipole expansion, resulting in significant rank reduction and signal loss, does not account for the adverse impact of the SSS filter on MVPA results.

A more plausible explanation for the detrimental effect of the SSS filter on MVPA is that the increase in white noise (random sensor noise) seen in Fig. 2 resulting from SSS filtering negatively impacts the performance of the SVM classifier. It is shown in the studies (Taulu and Kajola, 2005; Zhdanov et al., 2023) that the power of noise increase induced by the SSS filter depends on the number of sensors in the array; specifically, the more sensors there are, the less the increase in noise. This correlation aligns with the observation that SSS has a more pronounced effect on MVPA results from the 102-sensor magnetometer array (Fig. 3A) compared to the 204-sensor gradiometer array (Fig. 4A), and it does not have a significant adverse effect when applied to all 306 sensors combined (Fig. 4B). We computed a decoding curve for SSS-filtered data (*L*_*in*_ = 8, *L*_*out*_ = 3) that was low-pass filtered at 35 Hz (not shown here), which showed no discernible difference from the decoding curve without the 35 Hz low-pass filter. Therefore, the white noise increases over the entire spectrum including lower frequencies.

In summary, to optimize noise reduction in MVPA for OPM-MEG systems, we recommend considering methods such as SSP and HFC. HFC would be more beneficial for the experiments with a moving sensor array since it is a model-based approach denoising data in real-time (Tierney et al., 2022), whereas SSP assumes a stationary sensor array relative to the environment. Other approaches such as 3rd-order gradient noise reduction might also be efficient for both still and moving experiments (Vrba and Robinson, 2001). Note that 3rd-order gradient noise reduction requires a reference sensors array.

There are ongoing efforts to enhance SSS filters. The effectiveness of the SSS method relies on adequate spatial sampling of the magnetic field. This entails not only having a satisfactory density of sensors but also ensuring that the sensor array can accurately detect various field components (Taulu and Kajola, 2005; Nurminen et al., 2010). Before we were discussing the magnetometer or the planar gradiometer arrays, both measure only radial field direction. Gradiometers specifically measure the difference of the radial field across two points, capturing the spatial planar gradient. When using triaxial OPMs (Boto et al., 2022), each sensor measures the three orthogonal directions of the magnetic field. As such, applying a SSS filter to a triaxial sensor array, may be more optimal in terms of attenuation external noise sources and result only in a modest increase in white noise (Nurminen et al., 2013; Tierney et al., 2022) and thus not have a negative effect on MVPA classification accuracy. Another solution to mitigate the white noise increase induced by the SSS filter is to use numerical regularisation when calculating the weights for the multipole expansion presented (Holmes et al., 2023; Wang et al., 2023). Lastly, the HFC method improved using prolate spheroidal coordinates (Tierney et al., 2023) might be advantageous for the MVPA analysis since it lowers the variance of the data without suppressing any brain signal as opposed to currently used HFC and SSP methods.

In this work we also demonstrate that raw data from 102 gradiometers performs better than 102 magnetometers in terms of MVPA performance (Fig. 5A). Consequently, we argue that noise reduction by using synthetic gradiometers or the newly developed intrinsic optically pumped magnetic gradiometers (OPMG) (Cook et al., 2024) attenuates environmental noise and will improve MVPA performance. Additionally, Fig. 5B illustrates that choosing between magnetometers and planar gradiometers does not seem to be a design consideration when optimizing an MEG system for MVPA when denoising filters like SSP have been applied.

We showed that even with a relatively small number of magnetometers (30) in the array, the MVPA classification accuracy reaches saturation. Since we anticipate that OPM sensors will offer improved spatial resolution in capturing brain signals, this finding cannot be directly extrapolated to the OPM system. The OPM system might result in a higher total classification rate and benefit from a greater number of sensors to achieve its maximum performance potential (Wens, 2023). Nevertheless, it is reasonable to suggest that performance comparable to the SQUID system can be achieved with at least 30 sensors in the OPM array. It is important to note that the optimal number of 30 sensors is specific to the experimental paradigm used. For MVPA applied to tasks involving other modalities or tasks, a different number of sensors may be optimal.

In conclusion, MVPA has emerged as a powerful tool in cognitive neuroscience for detecting representational-specific neural patterns using M/EEG. Its application to OPM-MEG data holds great promise, particularly with the improved spatial coverage provided by the new system and the sensors being close to the head. Here we investigated the pre-processing pipeline when utilizing OPM-MEG data for MVPA. We recommend using filtering methods like HFC or SSP that reduce the noise floor of the data. We do not recommend using an SSS filter for MVPA with less than 300 sensors in the array.

## CRediT authorship contribution statement

Yulia Bezsudnova: Conceptualization; Formal analysis; Investigation; Methodology; Project administration; Visualization; Writing—Original draft; Writing—Review & editing. Andrew Quinn: Statistical analysis; Conceptualization. Ole Jensen: Conceptualization; Supervision; Funding Acquisition; Writing—Review & editing.

## Data Availability

Raw data is available upon request and code can be accessed on https://github.com/Y-Bezs/cross-modal-project

